# Differences in durability of PARP inhibition by clinically approved PARP inhibitors: implications for combinations and scheduling

**DOI:** 10.1101/2022.01.24.477471

**Authors:** Hannah L Smith, Elaine Willmore, Asima Mukhopadhyay, Yvette Drew, Nicola J Curtin

## Abstract

Five PARP inhibitors (PARPi) are approved for cancer treatment, they exploit cancer-specific defects in homologous recombination repair (HRR) to selectively kill tumour cells. Continuous PARP inhibition is required for single-agent anticancer activity. PARPi are also being investigated with ATR inhibitors clinically. We previously showed rucaparib caused prolonged PARP inhibition. Here we aimed to determine if this property was unique to rucaparib or common to other PARPis and the implications for scheduling with an ATR inhibitor (VE-821). Durability of PARP inhibition was determined at 0, 1, 24, 48 and 72 h after a 1 h pulse of 1μM of rucaparib, olaparib, niraparib, talazoparib or pamiparib in IGROV-1 (human ovarian cancer) cells. Inhibition of PARP was sustained to a variable degree with all inhibitors, but reduced with time. Rucaparib caused the most persistent inhibition of PARP activity, which was maintained at ≥75% for 72 h after drug withdrawal. In contrast, only 12% inhibition remained at this time with talazoparib and pamiparib and no detectable inhibition with olaparib and niraparib. Rucaparib enhanced VE-821 cytotoxicity to a similar extent in a sequential schedule as in co-exposure studies (PF_50_: 2.6 vs. 2.7) and there was even an approx. 2-fold enhancement after a 24 h delay between rucaparib and VE-821. Olaparib and niraparib produced similar enhancement of VE-821 cytotoxicity if co-exposed but were ineffective in sequential exposures. These data have clinical implications for both schedules of current PARPi monotherapy and the scheduling of PARPi in combination with ATRi and other cytotoxic drugs.

**Novelty and Impact:** PARPi are a new class of anticancer agent. We demonstrate for the first time that 5 PARPi continue to suppress cellular PARP activity after drug removal to a variable extent. Rucaparib caused the most durable PARP inhibition, olaparib and niraparib the least. Rucaparib enhanced ATR inhibitor cytotoxicity in sequential and co-exposures, olaparib and niraparib were only active in co-exposure settings. These data have implications for the clinical use of PARPi, particularly in combination with other drugs.

## Introduction

PARP inhibitors (PARPi) are a new class of anticancer drug that work by both inhibiting the repair of DNA single-strand breaks and trapping PARP1 at the site of the break ^1,2^. This results in replication-associated lesions that both activate ATR and the DNA damage cell cycle checkpoint cascade and which are resolved by homologous recombination DNA repair (HRR: Figure 1) ^3–5^.

**Figure 1:**
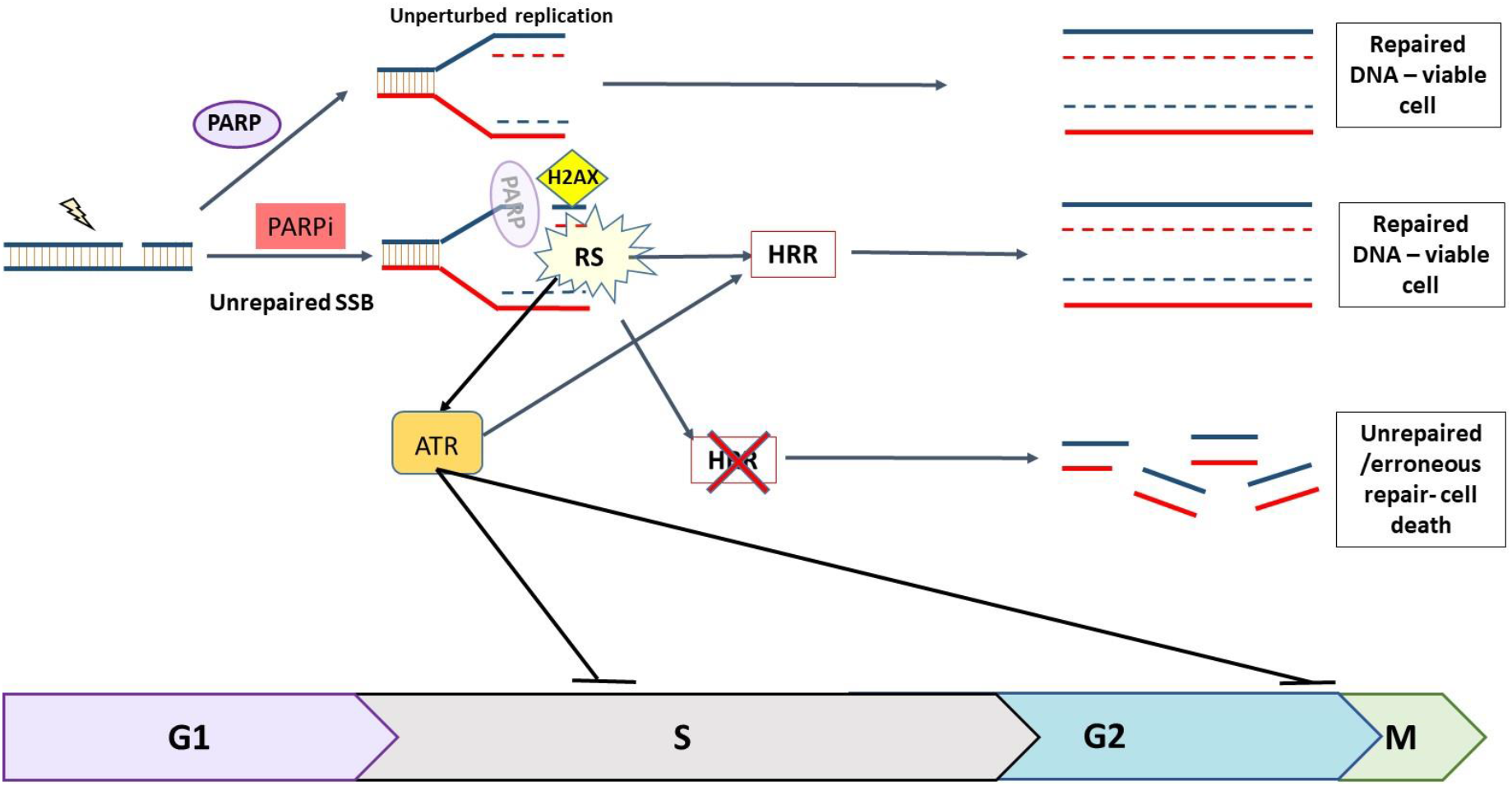
The role of PARP and ATR in the DNA damage response. Endogenously generated SSBs are continuously repaired by PARP-dependent repair mechanisms. When PARP is inhibited unrepaired SSB collide with replication forks causing them to stall and collapse resulting in DSBs which can only be repaired by HRR during S and early G2 phase. If HRR is defective, e.g. due to *BRCA* mutation the DNA cannot be repaired accurately, resulting in cell death. Replication stress (RS) caused by PARPi activates ATR which triggers a cascade which halts cell cycle progression and promotes HRR.

PARPi exploit tumour-specific defects in HRR, e.g. *BRCA* mutations, by a process known as synthetic lethality ^6–8^. The first approvals for PARPi by the FDA started in 2014 with olaparib and subsequently rucaparib, niraparib and talazoparib have been approved (https://www.fda.gov) and subsequently by the European Medicines Agency. Pamiparib was given conditional approval by China’s National Medical Products Administration (NMPA; https://www.pharmaceutical-technology.com/news/beigene-ovarian-cancer-drug/) in May 2O21.They are approved as single agents for cancers associated with defective HRR (HRD): ovarian, breast, castrate-resistant prostate cancer and pancreatic cancer. These and other PARPi are in advanced clinical trial and numerous trials are investigating combinations with cytotoxic and molecularly targeted therapy, including ATR inhibitors (https://clinicaltrials.gov). All approved PARPi are given continuously, on a daily or twice daily basis because, for effective single agent activity, PARP must be completely and continuously inhibited, so that cells cannot repair DNA breaks before S-phase progression ^9–11^. However, continuous inhibition may not be appropriate for combinations with genotoxic agents. It is becoming apparent that differences between the PARPi exist, in terms of their specificity, potency and “trapping” ability, which may underlie some of the differences in their activity and toxicity clinically ^12–15^. To date no study has investigated potential differences in the durability of PARP inhibition between the approved PARPi. Our previous studies indicated that rucaparib induced durable PARP inhibition in patients ^16^ and that a weekly schedule was as effective as daily dosing against *BRCA2* mutant xenografts^17^.

The purpose of the work reported here was to determine if persistent PARP inhibition was a class effect or unique to rucaparib using a GCLP-validated cell-based PARP activity assay. We used the clinically approved PARPi: olaparib, rucaparib, niraparib, talazoparib, and pamiparib. Pamiparib was of particular interest because it resembles rucaparib structurally in that the carboxamide group of the nicotinamide pharmacophore is incorporated into a 7-membered ring (Figure 2).

**Figure 2:**
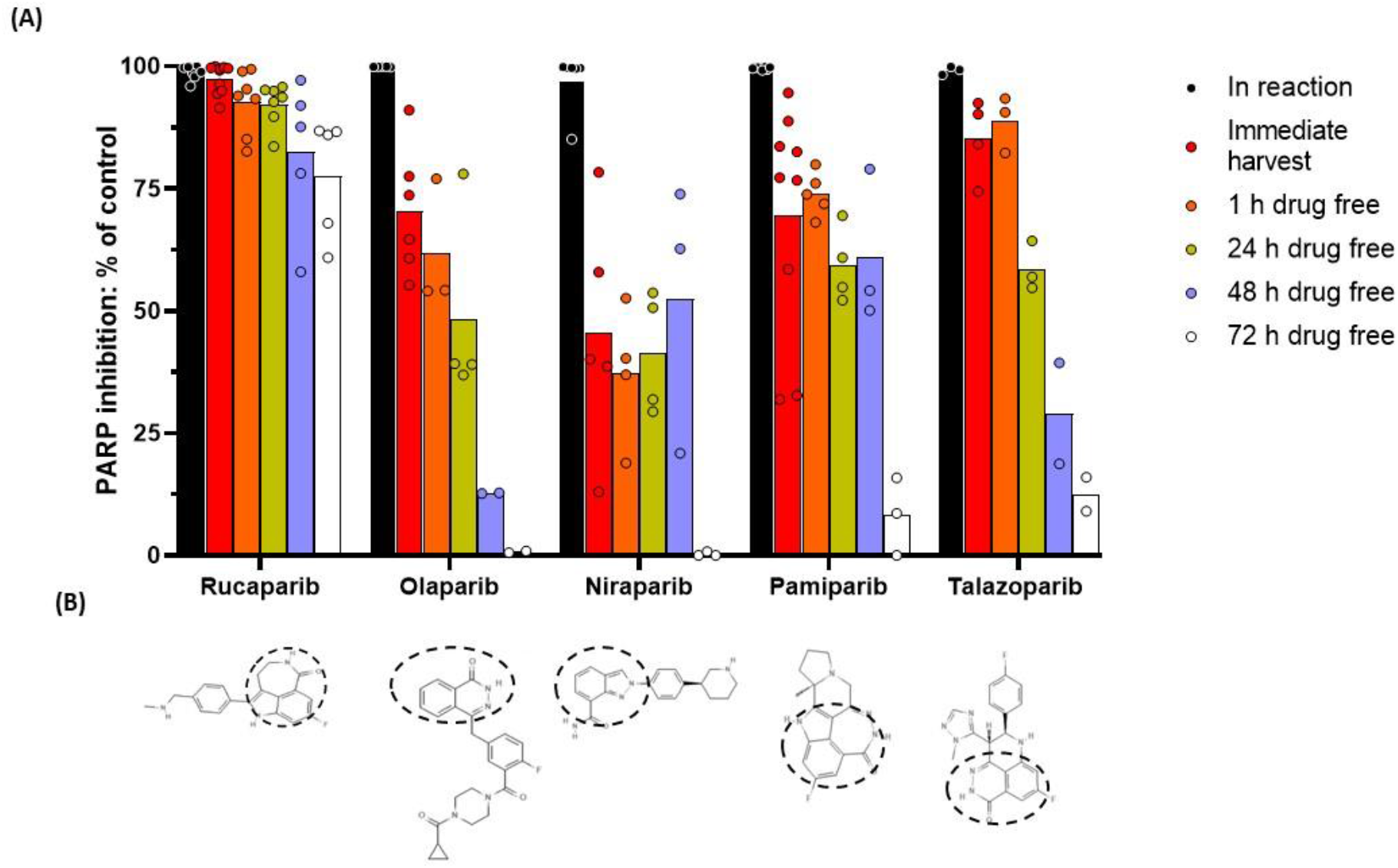
Durability of PARP activity inhibition by PARP inhibitors in IGROV-1 cells. **A.** PARP activity was measured in cells treated for 1 h with PARPi and harvested immediately and following 1, 24, 48 or 72 h in fresh media. Percent inhibition was calculated by comparison with untreated cells. Each data point on the graph represents an individual experiment and bars represent the mean of these experiments. **B.** Chemical structure of PARPis. Dashed circle highlights the nicotinamide pharmacophore in the PARPi structures.

The impact of scheduling on the synergy with ATR inhibitors was also investigated. We discovered that rucaparib was unique in its durability of target inhibition and was effective in sequential administration with the ATR inhibitor, VE-821. These data have clinical implications.

## Materials and Methods

### Chemicals and Reagents

The PARP inhibitor, rucaparib, was kindly gifted from Pfizer Global R&D and the ATR inhibitor, VE-821, was generously supplied by Merck (Merck KGaA, Darmstadt, Germany). Olaparib, niraparib, talazoparib and pamiparib were purchased from Selleckchem (Houston, TX, USA). Drugs dissolved in dry DMSO were stored at – 80 °C. Routine chemicals were of the highest purity from Sigma-Aldrich (Poole, United Kingdom), unless otherwise stated.

### Cell culture

IGROV-1 cells were obtained from American Type Culture Collection (ATCC; Manassas, VA, United States) and used within 30 passages of purchase or subsequent authentication by STR profiling (LGC Standards). They were maintained in exponential phase in RPMI-1640 medium (Merck, Kenilworth, NJ, United States) and 10% foetal bovine serum (FBS; Gibco, Life Technologies, CA, United States), at 37°C, 5% CO_2_ and 95% humidity. Cells were mycoplasma free.

### PARP activity assay

A GCLP-validated assay was used to measure DNA damage-activated PARP activity in permeabilised cells, in the presence of NAD+ substrate (350 nM) and a 12 mer palindromic double-stranded oligonucleotide (10 mg/ml) (Invitrogen, Waltham, MA, USA) to activate PARP1, by immunological detection of the product (PAR), using 10H Ab (Enzo life sciences, Farmingdale, NY, USA) and secondary HRP-conjugated goat anti-mouse Ab (Dako, Santa Clara, CA, USA), as described previously ^16–18^. IGROV-1 cells were exposed to 1 μM rucaparib, olaparib, niraparib, pamiparib or talazoparib for 1 h before drug was washed off and replaced with fresh media prior to cells being harvested immediately or after 1, 24, 48 or 72 h incubation in drug-free medium. Cells were permeabilised and PARP activity measured in comparison to untreated cells and cells with 1 μM drug added directly to the reaction mixture. The percentage PARP activity was calculated, relative to untreated cells.

### Cytotoxicity assays

Exponentially growing cells were seeded at various densities estimated to give 20-200 colonies following drug treatment. Cells were exposed to 0.5% DMSO (control) or VE-821 (1, 3 and 10 μM) or PARPi (1 μM) single agent in 0.5% DMSO for 24 h. Co-exposed cells were treated with VE-821 (1, 3 or 10 μM) and 1 μM PARPi for 24 h, before drug was removed and replaced with fresh media for colony formation. Sequentially exposed cells were treated with PARPi for 24 h before replacement with media containing VE-821 for another 24 h then drug-free medium for colony formation. Delayed sequentially exposed cells were similarly treated with PARPi for 24 h but media was replaced with fresh drug-free media for a further 24 h prior to 24 h exposure to VE-821 then drug-free medium for colony formation media. Colonies were fixed after 10-14 days in methanol: acetic acid (3:1) and stained with 0.4% crystal violet before colonies of >30 cells were counted by eye. Cell survival was calculated from the number of colonies relative to the number of cells seeded. Data were normalised to DMSO control or rucaparib alone, as appropriate, and plotted using Graphpad Prism 9.0 software (San Diego, CA, USA). The potentiation factor, PF_50_ is a unitless variable calculated as the lethal concentration 50% (LC_50_) for VE-821 alone/LC_50_ VE-821 + PARPi. Statistical analyses were performed using GraphPad Prism 9.0.

## Results

### Durability of PARPi by rucaparib, olaparib, niraparib, pamiparib and talazoparib

We first investigated whether our previously observed persistent PARP inhibition by rucaparib was unique or a class effect by comparing the durability of PARP inhibition by rucaparib, olaparib, niraparib, pamiparib and talazoparib in ovarian IGROV-1 cells. These cells were selected as previous studies with rucaparib have been done in SW620, Capan-1 and MX-1 cells, with similar results observed in all 3 cell lines ^17^. We therefore believe that inhibition of PARP activity is not cell line dependent. PARPi are currently used most often in the treatment of ovarian cancer, so an ovarian cancer cell line was selected for this study.

All of the PARPi inhibited PARP activity completely at 1 μM when added directly into the reaction mix (Figure 2). However, this level of PARP inhibition in the cells harvested immediately after 1 h exposure to PARPi was only observed with rucaparib with the other PARPi having reduced levels of inhibition in the range of 47% (niraparib) to 85% (talazoparib). This could be due to failure to achieve sufficient intracellular concentration or wash-out during the harvesting process, as similar levels of inhibition were noted after a further 1 h in drug-free medium. The extent of PARP inhibition decreased over time with all inhibitors but the rate of decrease differed substantially between the inhibitors. The decline in inhibition was slowest with rucaparib with >90% inhibition for 24 h and fastest with olaparib and niraparib. Notably, after 72 h in drug-free medium, rucaparib-treated cells still retained 75% PARP inhibition whereas there was only 12 % inhibition for pamiparib and talazoparib, and undetectable inhibition after exposure to olaparib and niraparib.

### Investigating schedules of PARPi and ATRi

To determine if the difference in durability of target inhibition by the PARPi would affect the cytotoxicity in combination with VE-821 on different schedules we compared the effect of PARPi on VE-821 cytotoxicity using 3 different schedules: concurrent, sequential PARPi followed by VE-821 and sequential with a 24 h drug-free gap between PARPi and VE-821 exposure. Rucaparib, the most durable, and olaparib and niraparib, as the least durable, were selected for this investigation (Figure 3).

**Figure 3:**
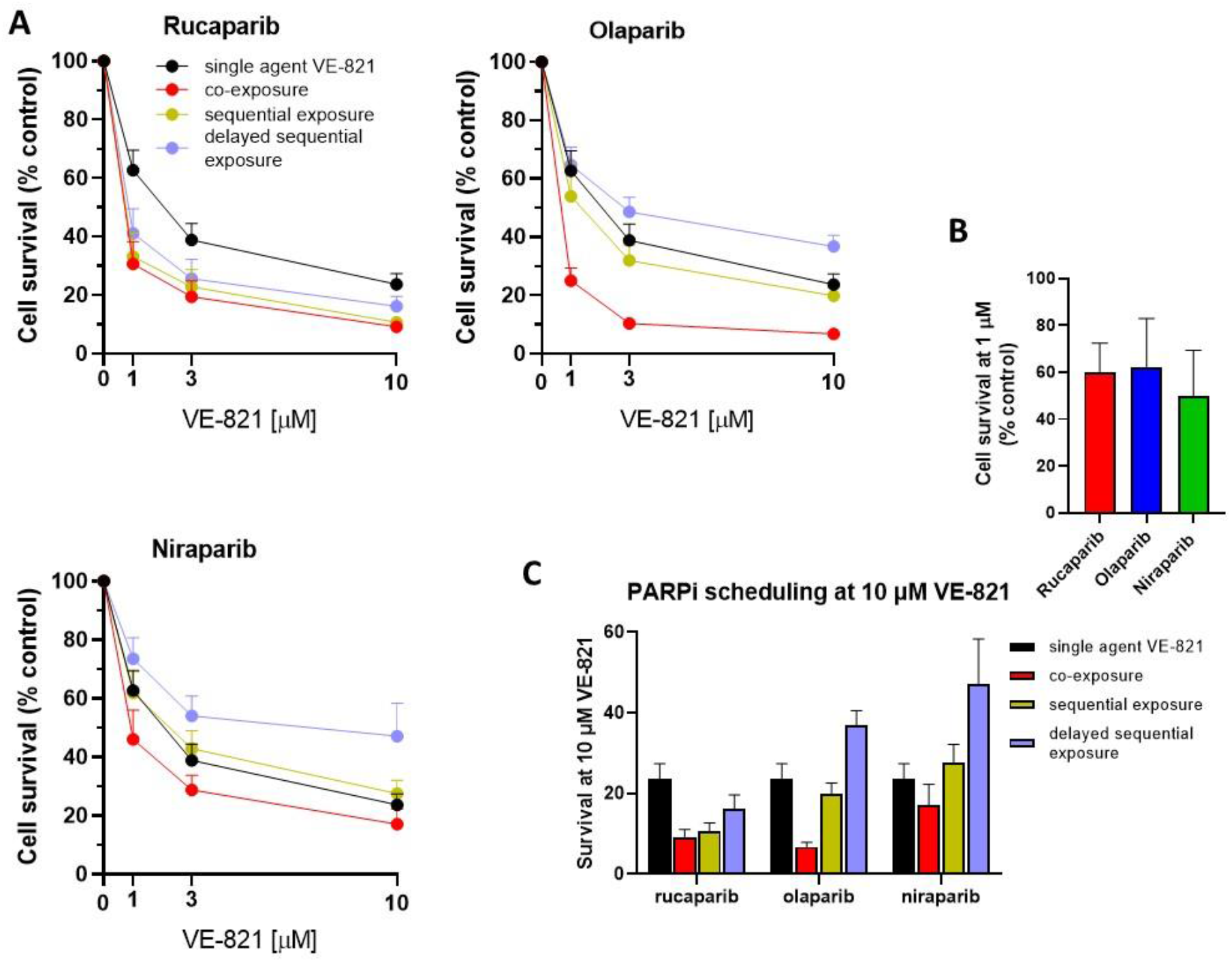
Scheduling exposure to PARPi and VE-821 in IGROV-1 cells. **A.** Cells were exposed to VE-821 at indicated concentrations either as single agent or with 1 μM rucaparib, olaparib or niraparib, either as co-exposure, sequential exposure or 24 h delayed sequential exposure then incubated for 10-14 days for colony formation. **B.** Cells were exposed to 1 μM rucaparib, olaparib and niraparib as single agents for 24 h then drug-free medium cells for 10-14 days and cell survival calculated by reference to vehicle alone (DMSO) controls, data are mean ± S.D of 5 independent experiments. **C.** Cell survival following exposure to 10 μM VE-821 alone or with 1 μM rucaparib, olaparib or niraparib, either as co-exposure, sequential exposure or 24 h delayed sequential exposure Data are representative of the mean and standard error of 5 independent experiments.

The cytotoxicity of combinations with VE-821 were normalised to PARPi alone and the cytotoxicity of VE-821 alone was normalised to DMSO control. Rucaparib, olaparib and niraparib all sensitised IGROV-1 cells to VE-821 in co-exposure experiments (Figure 3A). Rucaparib and olaparib caused similar levels of sensitisation at the LC_50_ (PF_50_ values 2.7 vs. 2.8, respectively) and at 10 μM VE-821 (3.5 vs. 3.8-fold, respectively), whereas niraparib had less impact sensitising cells when co-exposed with VE-821 (PF_50_: 1.8 and sensitisation at 10 μM: 1.7-fold). This was not due to any difference in the intrinsic cytotoxicity of the PARPi alone as there was no significant difference between the inhibitors, in terms of cytotoxicity with 1 μM causing between 40 and 50% inhibition of cell viability (Figure 3B). With sequential exposure, where cells were incubated with the PARPi alone for 24 h immediately followed by 24 h exposure to VE-821 alone, rucaparib was equally as effective as when the exposure with VE-821 was concurrent (PF_50_: 2.6, potentiation at 10 μM: 2.5-fold), indicating PARP inhibition was sustained for 24 h (Figure 3A). In marked contrast, neither niraparib nor olaparib significantly increased VE-821 cytotoxicity when given sequentially.

When the cells were incubated for 24 h in drug-free medium between the PARPi and VE-821 exposures rucaparib still caused an approximately 2-fold sensitisation at the LC_50_ and 10 μM VE-821 (2.2 and 1.8-fold, respectively) indicating some PARP inhibition is sustained over the 24-48 h period after rucaparib withdrawal. No potentiation was observed with olaparib or niraparib, in fact there seemed to be some modest protection (Figure 3A and C). This observed protection was significant at 10 μM VE-821 with olaparib (p= 0.005) (Figure 3C), which may indicate olaparib exposure had synchronised the cells such that they were not in S-phase during incubation with VE-821. This protection was not significant in cells treated with niraparib.

## Discussion

Following previous studies highlighting the durability of PARP inhibition by rucaparib ^16,17^ we investigated if this was a unique property of this PARPi or was common to other clinically active PARPi. Our results reveal that although all PARPi continued to inhibit cellular PARP activity to a certain extent after drug removal, the sustained inhibition of PARP activity over a period of 3 days was unique to rucaparib. There was a progressive loss of PARP inhibition between 24 and 72 h post drug removal with all inhibitors, which was most rapid for olaparib and niraparib and intermediate for talazoparib and pamiparib. Only for rucaparib was inhibition ≥75% for 72 h. Notably, pamiparib did not have similar levels of durability observed in rucaparib, despite some structural similarities. Our previous studies indicated that rucaparib uptake into cells was carrier-mediated and that it was retained ^17^, it has not been possible to repeat such studies with the other PARPi as radiolabelled compound is not available. These results also suggest durability is not related to trapping potency as talazoparib is the most potent trapper ^1^ but clearly not the most durable.

For single agent activity PARP must be inhibited continuously and all PARPi are approved for daily or twice daily administration. The approved doses and schedules of the PARPi are established in early phase trials based on PK and tolerability rather than optimum pharmacodynamic effect. Rucaparib is recommended at a dose of 600 mg twice daily but its unique ability to cause sustained PARP inhibition suggests that twice daily dosing may not be necessary and an intermittent dosing schedule, perhaps twice weekly, might be equally effective, as well as more tolerable and affordable. Indeed, our earlier xenograft studies would suggest that this is plausible ^17^ and clinical evaluation of intermittent schedules is warranted. Similarly, talazoparib appears to cause >80% PARP inhibition for 24 h and could potentially be given on alternate days. A reduced dose-intense schedule would be particularly beneficial in low to middle-income countries where for the majority of women the current recommended dose is unaffordable.

Following sound preclinical evidence demonstrating the synergy between PARPi and ATRi ^19–22,3^ PARPi are currently being investigated clinically in combination with ATR inhibitors. Rucaparib is not currently being investigated in combination with ATR inhibitors in the clinical setting. However, olaparib is in various Phase 2 trials with the ATR inhibitor, AZD6738, for a variety of cancer types (NCT03682289, NCT03787680, NCT03462342, NCT04065269) and niraparib is in Phase 1 trials in combination with various ATR inhibitors (NCT03682289, NCT03787680, NCT03462342, NCT04065269, NCT04170153, NCT04149145, NCT04267939). Our investigation of various schedules of administration of PARPi with the ATR inhibitor VE-821 revealed that olaparib and niraparib only synergised with VE-821 when cells were exposed to both drugs concurrently. In contrast, rucaparib was similarly effective in enhancing VE-821-mediated cytotoxicity when given simultaneously or immediately prior to VE-821 exposure and even increased the cytotoxicity of VE-821 approximately 2-fold when there was a 24 h delay in adding the VE-821. On the basis of these data, it may not be appropriate to extrapolate from trials with olaparib and niraparib when designing rucaparib combination trials. Furthermore, it may be possible to consider schedules where rucaparib is given on an intermittent basis (every 2 or 3 days) in combination with an ATRi. Such schedules may be equally effective but less toxic. However, further investigations are required to determine whether sequential dosing would have equivalent anti-cancer activity *in vivo* and to examine the implications for toxicity prior to exploring different schedules clinically.

The toxicities associated with PARPi and cytotoxic chemo- and radiotherapy clinically have resulted in a failure of PARPi to progress beyond Phase 2 clinical evaluation in combination with cytotoxic therapy, even though this was the original purpose of PARPi ^16,23^. Preclinical data indicates that for synergy with cytotoxic chemo- and radiotherapy lower doses and shorter duration of PARP inhibition, i.e., during the repair phase of the induced damage, is all that is necessary and that higher doses are highly toxic ^23,24^. The data reported here have implications for the investigation of such combinations clinically. It could be that the modest persistence in PARP inhibition after removal of drug with all of the inhibitors, not just rucaparib, accounts for some of the clinical toxicities observed and that shorter PARPi schedules would be preferable.

In conclusion, we report here that 5 clinically active PARPi continue to inhibit cellular PARP activity after the drug has been removed but this inhibition diminishes with time to a variable degree. Suppression of PARP activity is most durable with rucaparib and least durable with olaparib and niraparib. This durable suppression meant that rucaparib was effective in enhancing ATR inhibitor-induced cytotoxicity when given prior to, including with a 24 h delay, ATR inhibition. In contrast, olaparib and niraparib only increased ATR inhibitor-induced cytotoxicity when given simultaneously. These data have implications for the scheduling of PARPis alone and in combination with ATR inhibitors and cytotoxic drugs.

## Acknowlegements, author contributions and conflict of interest

H.L.Smith was supported in conducting this study partially by seedcorn funding from CRUK/Wellcome Trust/Indian Department of Technology Affordable approaches to cancer and NJC’s internal funding. NJC, AM and YD conceived the study, HLS and NJC wrote the manuscript which was edited by YD and EW. We are grateful to Frank Zenke (Merk) and Thomas Harding (Clovis) for helpful comments. NJC, YD and AM were involved in the development of rucaparib and have received research funding and royalty payments as a result. NJC and YD have also received research funding from Merck. All other authors have no conflict of interest.

## Statement of ethics

Ethical approval for the Newcastle Academic Health Partners Bioresource was obtained from North East Newcastle and North Tyneside Research Ethics Committee 1 for the collection of clinical material and patient data (REC 17/NE/0361, REC 12/NE/0395). All patients gave written informed consent and samples were anonymised.

## Abbreviations

ATR: Ataxia telangiectasia and Rad3 related kinase
FDA: US Food and Drug Administration
GCLP: Good Clinical Laboratory Practice
HRR: Homologous recombination DNA repair
PARP: poly(ADP-ribose) polymerase
LC50: Lethal concentration 50, the concentration killing 50% of cells
PF_50_: Potentiation factor (fold potentiation) at the LC50

## References

1 Murai, J., Huang, S.Y., Das, B.B., Renaud, A., Zhang, Y., Doroshow, J.H., Ji, J., Takeda, S. and Pommier, Y. Trapping of PARP1 and PARP2 by Clinical PARP Inhibitors. Cancer Res 2012; 72(21): p. 5588–99.

2 Rose, M., Burgess, J.T., O’Byrne, K., Richard, D.J. and Bolderson, E. PARP Inhibitors: Clinical Relevance, Mechanisms of Action and Tumor Resistance. Front Cell Dev Biol 2020; 8: p. 564–601.

3 Gralewska, P., Gajek, A., Marczak, A., Mikuła, M., Ostrowski, J., Śliwińska, A. and Rogalska, A. PARP Inhibition Increases the Reliance on ATR/CHK1 Checkpoint Signaling Leading to Synthetic Lethality-An Alternative Treatment Strategy for Epithelial Ovarian Cancer Cells Independent from HR Effectiveness. Int J Mol Sci 2020; 21(24).

4 Yazinski, S.A., Comaills, V., Buisson, R., Genois, M.M., Nguyen, H.D., Ho, C.K., Todorova Kwan, T., Morris, R., Lauffer, S., Nussenzweig, A., Ramaswamy, S., Benes, C.H., Haber, D.A., Maheswaran, S., Birrer, M.J. and Zou, L. ATR inhibition disrupts rewired homologous recombination and fork protection pathways in PARP inhibitor-resistant BRCA-deficient cancer cells. Genes Dev 2017; 31(3): p. 318–332.

5 Smith, H.L., Southgate, H., Tweddle, D.A. and Curtin, N.J. DNA damage checkpoint kinases in cancer. Expert Rev Mol Med 2020; 22.

6 Bryant, H.E., Schultz, N., Thomas, H.D., Parker, K.M., Flower, D., Lopez, E., Kyle, S., Meuth, M., Curtin, N.J. and Helleday, T. Specific killing of BRCA2-deficient tumours with inhibitors of poly(ADP-ribose) polymerase. Nature 2005; 434(7035): p. 913–917.

7 Farmer, H., McCabe, N., Lord, C.J., Tutt, A.N., Johnson, D.A., Richardson, T.B., Santarosa, M., Dillon, K.J., Hickson, I., Knights, C., Martin, N.M., Jackson, S.P., Smith, G.C. and Ashworth, A. Targeting the DNA repair defect in BRCA mutant cells as a therapeutic strategy. Nature 2005; 434(7035): p. 917–921.

8 Setton, J., Zinda, M., Riaz, N., Durocher, D., Zimmermann, M., Koehler, M., Reis-Filho, J.S. and Powell, S.N. Synthetic Lethality in Cancer Therapeutics: The Next Generation. Cancer Discov 2021; 11(7): p. 1626–1635.

9 Javle, M. and Curtin, N.J. The role of PARP in DNA repair and its therapeutic exploitation. Br J Cancer 2011; 105(8): p. 1114–1122.

10 Michelena, J., Lezaja, A., Teloni, F., Schmid, T., Imhof, R. and Altmeyer, M. Analysis of PARP inhibitor toxicity by multidimensional fluorescence microscopy reveals mechanisms of sensitivity and resistance. Nat Commun 2018; 9(1): p. 2678.

11 Simoneau, A., Xiong, R. and Zou, L. The *trans* cell cycle effects of PARP inhibitors underlie their selectivity toward BRCA1/2-deficient cells. Genes Dev 2021; 35(17-18): p. 1271–1289.

12 Murthy, P. and Muggia, F. Women’s cancers: how the discovery of BRCA genes is driving current concepts of cancer biology and therapeutics. Cancer medical science 2019; 13: p. 904.

13 Dickson, K.A., Xie, T., Evenhuis, C., Ma, Y. and Marsh, D.J. PARP Inhibitors Display Differential Efficacy in Models of BRCA Mutant High-Grade Serous Ovarian Cancer. Int J Mol Sci 2021; 22(16).

14 Cai, Z., Liu, C., Chang, C., Shen, C., Yin, Y., Yin, X., Jiang, Z., Zhao, Z., Mu, M., Cao, D., Zhang, L. and Zhang, B. Comparative safety and tolerability of approved PARP inhibitors in cancer: A systematic review and network meta-analysis. Pharmacol Res 2021; 172: p. 105808.

15 Krastev, D.B., Wicks, A.J. and Lord, C.J. PARP Inhibitors - Trapped in a Toxic Love Affair. Cancer Res 2021; 81(22): p. 5605–5607.

16 Plummer R, Jones C, Middleton M, et al. Phase I study of the poly(ADP-ribose) polymerase inhibitor, AG014699, in combination with temozolomide in patients with advanced solid tumors. Clin Cancer Res. 2008;14(23):7917–7923.

17 Murray, J., Thomas, H., Berry, P., Kyle, S., Patterson, M., Jones, C., Los, G., Hostomsky, Z., Plummer, E.R., Boddy, A.V. and Curtin, N.J. Tumour cell retention of rucaparib, sustained PARP inhibition and efficacy of weekly as well as daily schedules. Br J Cancer 2014; 110(8): p. 1977–84.

18 Canan, S., Maegley, K. and Curtin, N.J. Strategies Employed for the Development of PARP Inhibitors. Methods Mol Biol 2017; 1608: p. 271–297.

19 Peasland, A., Wang, L.Z., Rowling, E., Kyle, S., Chen, T., Hopkins, A., Cliby, W.A., Sarkaria, J., Beale, G., Edmondson, R.J. and Curtin, N.J. Identification and evaluation of a potent novel ATR inhibitor, NU6027, in breast and ovarian cancer cell lines. Br J Cancer 2011; 105(3): p. 372–81.

20 Middleton, F.K., Patterson, M.J., Elstob, C.J., Fordham, S., Herriott, A., Wade, M.A., McCormick, A., Edmondson, R., May, F.E., Allan, J.M., Pollard, J.R. and Curtin, N.J. Common cancer-associated imbalances in the DNA damage response confer sensitivity to single agent ATR inhibition. Oncotarget 2015; 6(32): p. 32396–409.

21 Kim, H., Xu, H., George, E., Hallberg, D., Kumar, S., Jagannathan, V., Medvedev, S., Kinose, Y., Devins, K., Verma, P., Ly, K., Wang, Y., Greenberg, R.A., Schwartz, L., Johnson, N., Scharpf, R.B., Mills, G.B., Zhang, R., Velculescu, V.E., Brown, E.J. and Simpkins, F. Combining PARP with ATR inhibition overcomes PARP inhibitor and platinum resistance in ovarian cancer models. Nat Commun 2020; 11(1): p. 3726.

22 Lloyd, R.L., Wijnhoven, P.W.G., Ramos-Montoya, A., Wilson, Z., Illuzzi, G., Falenta, K., Jones, G.N., James, N., Chabbert, C.D., Stott, J., Dean, E., Lau, A. and Young, L.A. Combined PARP and ATR inhibition potentiates genome instability and cell death in ATM-deficient cancer cells. Oncogene 2020; 39(25): p. 4869–4883.

23 Curtin, N.J. and Szabo, C. Poly(ADP-ribose) polymerase inhibition: past, present and future. Nat Rev Drug Discov 2020; 19(10): p. 711–736.

24 Calabrese, C.R., Almassy, R., Barton, S., Batey, M.A., Calvert, A.H., Canan-Koch, S., Durkacz, B.W., Hostomsky, Z., Kumpf, R.A., Kyle, S., Li, J., Maegley, K., Newell, D.R., Notarianni, E., Stratford, I.J., Skalitzky, D., Thomas, H.D., Wang, L.Z., Webber, S.E., Williams, K.J. and Curtin, N.J. Anticancer chemosensitization and radiosensitization by the novel poly(ADP-ribose) polymerase-1 inhibitor AG14361. J Natl Cancer Inst 2004; 96(1): p. 56–67.

